# Nucleosomal association and altered interactome underlie the mechanism of cataract caused by the R54C mutation of αA-crystallin

**DOI:** 10.1101/2020.07.03.182295

**Authors:** Saad M. Ahsan, Bakthisaran Raman, Tangirala Ramakrishna, Ch. Mohan Rao

## Abstract

The small heat shock protein (sHSP), αA-crystallin, plays an important role in eye lens development. It has three distinct domains *viz*. the N-terminal domain, α-crystallin domain and the C-terminal extension. While the α-crystallin domain is conserved across the sHSP family, the N-terminal domain and the C-terminal extension are comparatively less conserved. Nevertheless, certain arginine residues in the N-terminal region of αA-crystallin are conserved across the sHSP family. Interestingly, most of the cataract-causing mutations in αA-crystallin occur in the conserved arginine residues. In order to understand the molecular basis of cataract caused by the R54C mutation in human αA-crystallin, we have compared the structure, chaperone activity, intracellular localization, effect on cell viability and “interactome” of wild-type and mutant αA-crystallin. Although R54CαA-crystallin exhibited slight changes in quaternary structure, its chaperone activity was comparable to that of the wild-type. When expressed in lens epithelial cells, R54CαA-crystallin triggered a stress-like response, resulting in nuclear translocation of αB-crystallin, disassembly of cytoskeletal elements and activation of Caspase 3, leading to apoptosis. Comparison of the “interactome” of the wild-type and mutant proteins revealed a striking increase in the interaction of the mutant protein with nucleosomal histones (H2A, H2B, H3 and H4). Using purified chromatin fractions, we show an increased association of R54CαA-crystallin with these nucleosomal histones, suggesting the potential role of the mutant in transcriptional modulation. Thus, the present study shows that alteration of “interactome” and its nucleosomal association, rather than loss of chaperone activity, is the molecular basis of cataract caused by the R54C mutation in αA-crystallin.

## 1 Introduction

αA- and αB-crystallin are members of the sHsp family and are abundantly present in the mammalian eye lens. They form a large multimeric complex, α-crystallin, in the mammalian eye lens, in which the two gene products *viz*. αA- and αB-crystallin are present in a 3:1 molar ratio [1, 2]. Both αA- and αB-crystallin have an oligomeric mass in the range of 600 to 800 kDa with monomer mass of about 20 kDa. They may assemble to form homo-oligomers and may form hetero-oligomers with other members [3]. These assemblies are dynamic in nature with a continuous exchange of subunits between the oligomers. These proteins have been demonstrated to exhibit molecular chaperone-like activity and are known to prevent the aggregation of other proteins [4–8], under normal [9–11] and stress conditions [12–15]. They work in an ATP-independent manner and prevent target protein aggregation, thereby keeping them in an active folded state.

The sHSPs are characterized by a well-conserved α-crystallin domain that is flanked by a comparatively less conserved N-terminal domain and a C-terminal extension. The N-terminal domain is known to be important in oligomerization, subunit exchange and chaperone-like activity [16]. Previous studies show that the chaperone-like activity of these proteins can be attributed to various regions spread over the entire length of the sHSP sequence [17]. Studies from our laboratory and others have reported the importance of the N-terminal domain, particularly the SRLFDQFFG stretch, in chaperone-like activity and oligomerization of alpha crystallins [17–19].

Many mutations that cause cataract have been reported in the N-terminal domain of αA-crystallin, both in humans and mice. Interestingly, most of these mutations involve the replacement of the conserved arginine residues at the 12^th^, 21^st^, 49^th^ and 54^th^ position **(figure 1)** [20–24]. However, the molecular basis of the mutation-induced cataract formation is not yet well understood. The chaperone-like activity was found to be compromised in several mutations. However, there are a few cases where the chaperone activity seems to be either unaltered or to be increased to some extent [25–27]. Therefore, it appears that factors other than the compromised chaperone activity and aggregation propensity also play a deleterious role, which is yet to be understood. For example, the R54C mutation in αA-crystallin leads to congenital cataract [23]. However, its chaperone activity is reported to be not altered significantly [25]. We, therefore, verified the structural and chaperone activity alteration, if any, caused by the R54C mutation in α-crystallin. We have investigated the intra-cellular expression/localization and compared the “interactome” of the wild type and mutant αA-crystallin to unravel the molecular basis of the mutation-induced pathology. Interestingly, our study reveals an increased interaction of the mutant with nucleosomal histones. This altered interaction may be a cause of a stress-like response associated with the mutant, leading to cytoskeleton disintegration and apoptosis.

**Figure 1:**
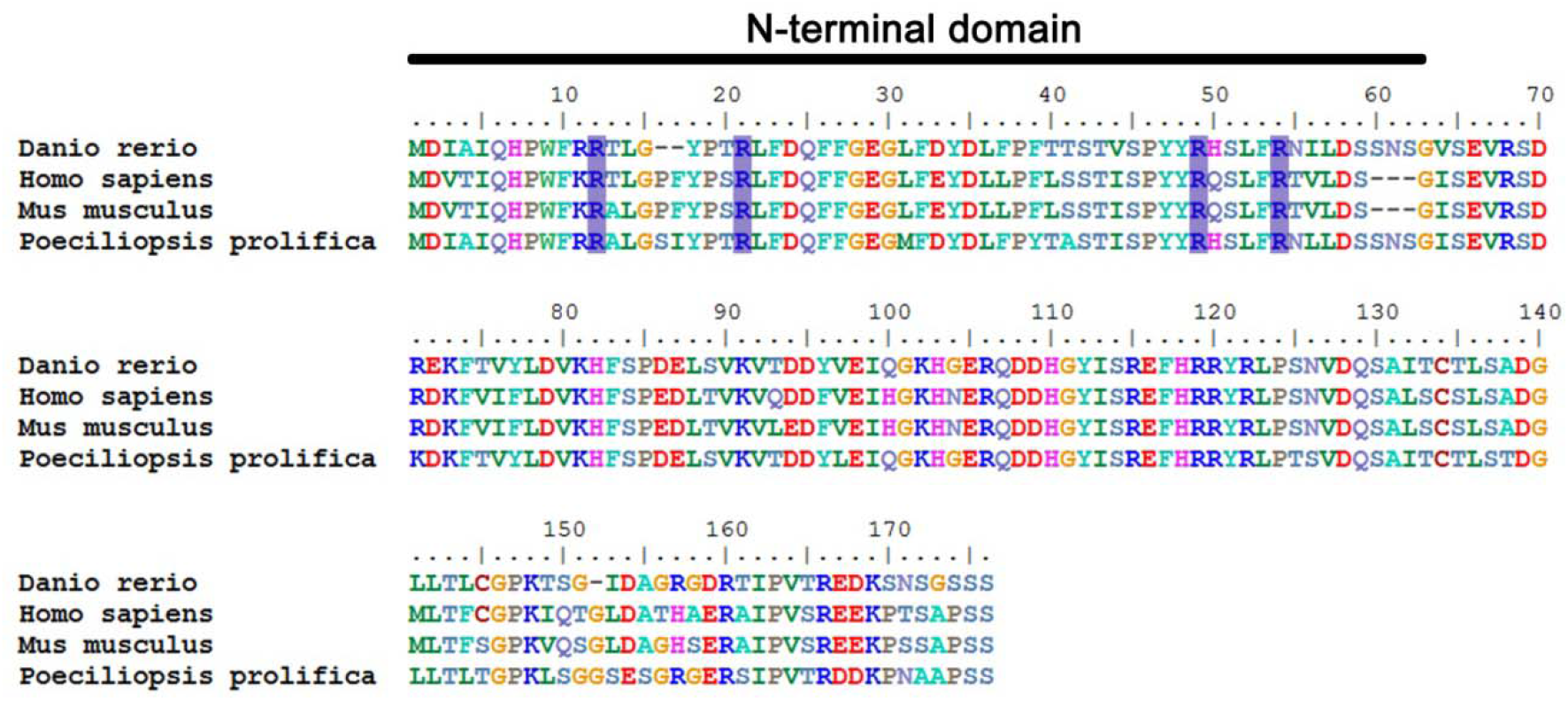
Sequence alignment of αA-crystallin from different species. The alignment reveals the conserved nature of the protein across different phyla. Highlighted in blue are the four well-conserved Arginine residues in the N-terminal domain, mutations in which lead to cataract.

## 2 Results

### 2.1 R54C mutation-induced structural alteration of αA-crystallin

To study the effect of the R54C mutation on the structure of human αA-crystallin, we expressed wild type and mutant recombinant proteins in *E.coli* BL21 (DE3). Like the wild type αA-crystallin, the mutant protein also partitioned exclusively in to the soluble fraction when expressed in the bacterial cells. The wild type and mutant proteins were purified using a method described for the purification of wild type αA-crystallin earlier [6] (briefly mentioned in section 2.3).

Far-UV circular dichroism (CD) spectroscopy was performed to compare the secondary structures of the wild type and the mutant protein. As shown in **figure 2A**, both the mutant (grey curve) and wild type (black curve) protein showed a minimum around 217 nm, indicative of a β-sheet structure. The mutant R54C, however, showed an increase in the negative ellipticity. A CAPITO (**C**D **A**nalysis and **P**lott**i**ng **To**ol) [28] analysis of the CD data indicates an increase in alpha-helical content **(table 1)**. The increased chirality (negative ellipticity) in this region may also be attributed to changes in dihedral angles of certain regions in the mutant sequence [29]. The near-UV CD spectra of mutant and wild type αA-crystallin show marginal changes **(figure 2B)**.

**Figure 2:**
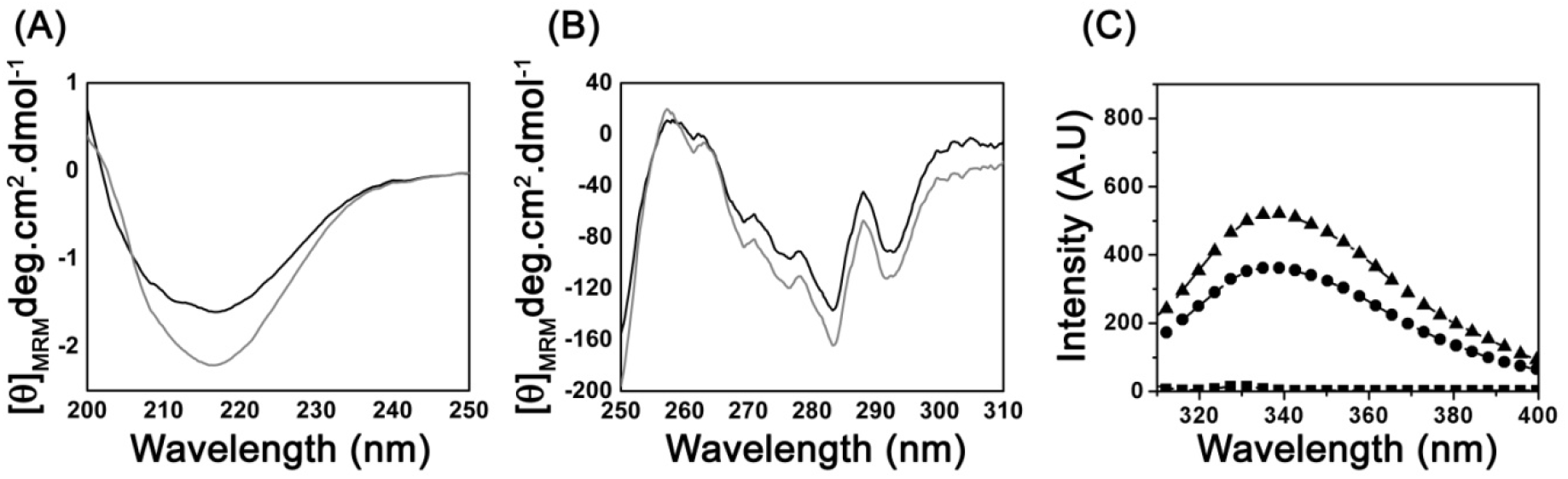
Secondary and tertiary structural studies on the wild type and R54C αA-crystallin. Far-UV CD spectra **(A)** and near-UV CD spectra **(B)** of wild type (black line) and R54C mutant (grey line). **(C)** Tryptophan fluorescence spectra of wild type (black circles) and R54C mutant (black triangles). Baseline buffer fluorescence is shown as black squares.

**Table 1:** Percent secondary structure content in wild type and R54C αA-crystallin:

| αA-crystallin | Helix | β-strand | Irregular |
| --- | --- | --- | --- |
| Wild type | 1.9 | 47.2 | 50.9 |
| R54C | 3.7 | 47.7 | 48.6 |

The intrinsic tryptophan fluorescence of the mutant and wild type proteins was studied in an attempt to gain insights into the microenvironment of the tryptophan (9th position) present in αA-crystallin. As the sole tryptophan in αA-crystallin is present in the N-terminal of the protein, it is expected to shed light on the folding of the N-terminal region of the protein. As shown in **figure 2C**, the tryptophan fluorescence intensity of the mutant was found to be increased slightly compared to the wild type protein, while the emission maximum remained the same (338 nm).

We investigated the quaternary structure/oligomeric size of the proteins by gel filtration chromatography using a Superose-6 column. Wild type αA-crystallin eluted out at a volume of 14.3 ml, whereas the R54C mutant eluted out at a slightly lower elution volume of 13.2 ml (**Figure 3**). The molecular mass of the wild type and mutant proteins, calculated using their elution volumes, were found to be 540 kDa and 720 kDa respectively. Dynamic light scattering study showed a slightly larger hydrodynamic radius for the mutant (9.3 nm) as compared to the wild type (8.7 nm). Overall, our structural studies indicate that the mutant protein exhibits a slightly altered secondary and tertiary structure and assembles into larger oligomers compared to the wild type αA-crystallin.

**Figure 3:**
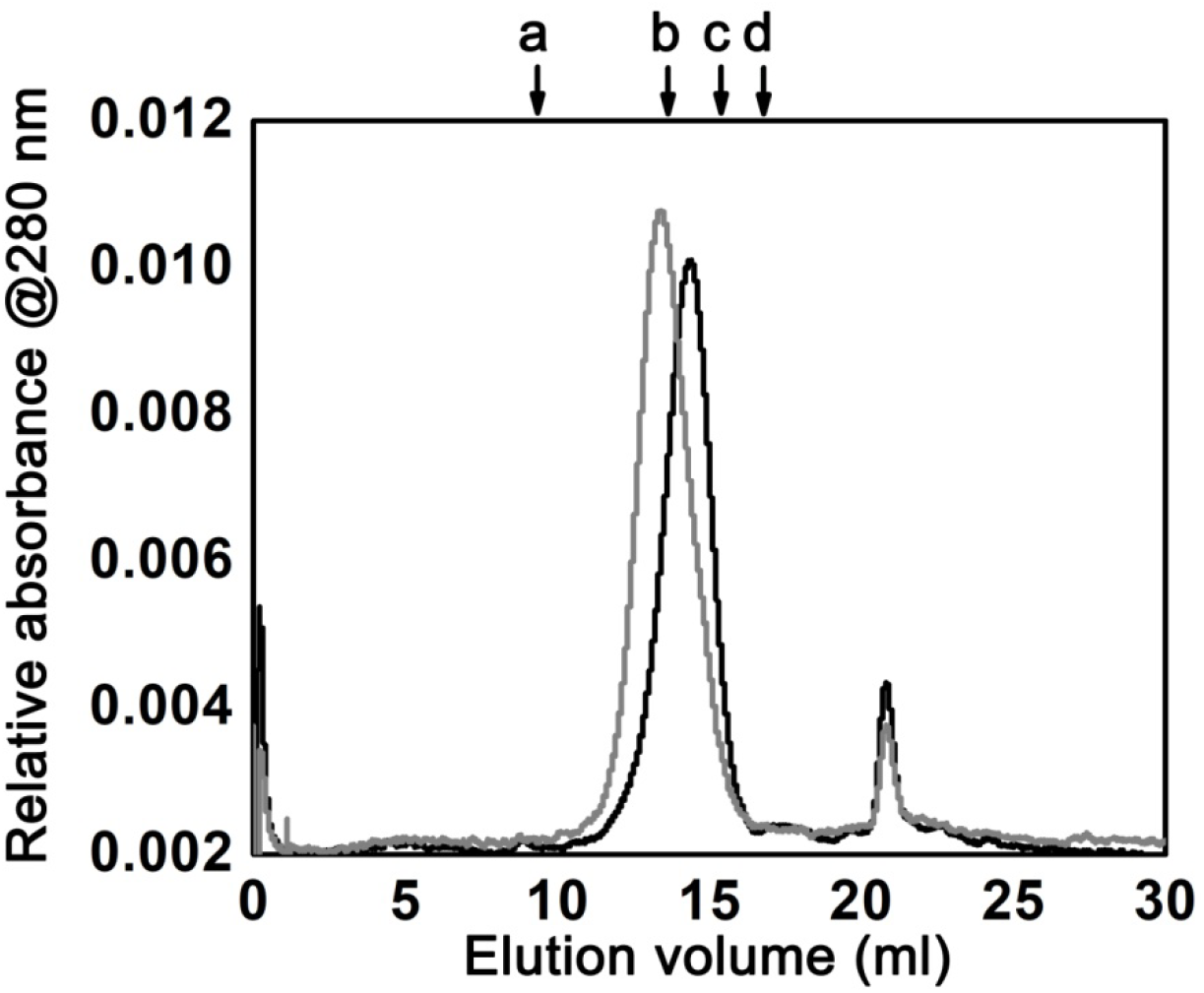
Quaternary structural analysis of wild type and R54C αA-crystallin. Elution profile of wild type (black line) and R54C mutant (grey line), as determined by size-exclusion chromatography. The retention volumes of the molecular mass standards are indicated by the arrows (a, blue dextran (2000 kDa); b, thyroglobulin (669 kDa); c, ferritin (440 kDa); d, catalase (232 kDa)). (Note: y-axis is expanded to highlight the difference in the peak positions)

### 2.2 Surface hydrophobicity

We probed the hydrophobic patches on the wild type and mutant αA-crystallin using bis-ANS (a hydrophobic probe). The fluorescence intensity of bis-ANS increases and its emission maximum shifts to lower wavelengths upon binding to hydrophobic surfaces of a protein [30]. In our experiments, the wild type and mutant αA-crystallin did not show any variations in emission maxima of bis-ANS fluorescence. The fluorescence intensity of bis-ANS was found to increase upon binding to wild type as well as R54C αA-crystallin, the increase being slightly higher for the mutant protein compared to that for the wild type protein (**figure 4A**).

The grand average of hydropathicity (GRAVY) index as determined by the ProtScale prediction suggested an increase in the local hydrophobicity around the substitution site (**figure 4B and 4C)**.

**Figure 4:**
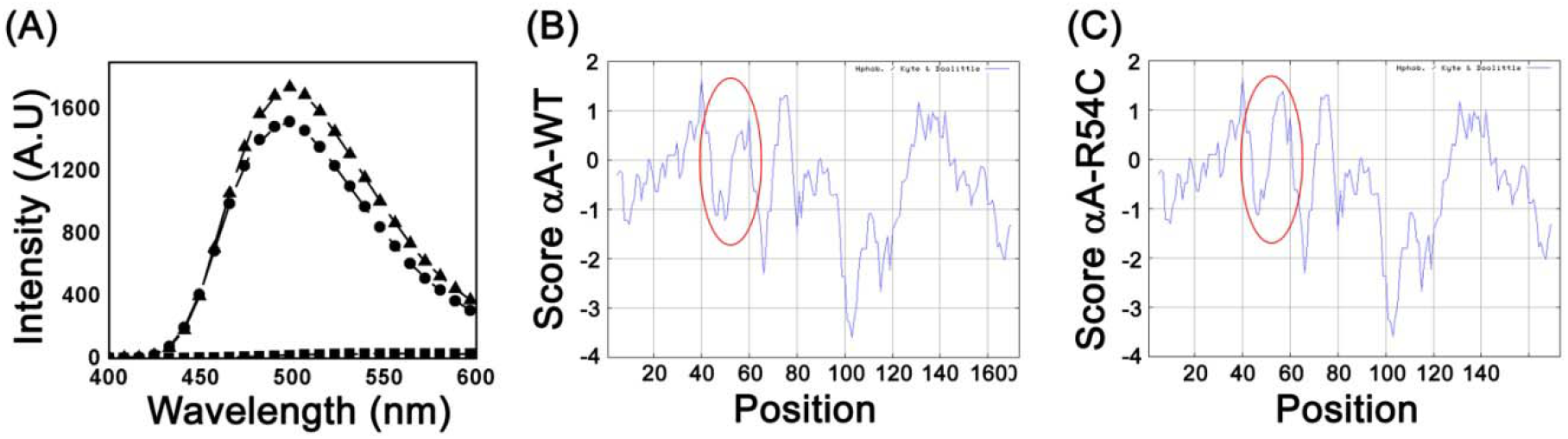
Hydrophobicity analysis of wild type and R54C αA-crystallin. **(A)** Fluorescence emission by bis-ANS upon binding to wild type αA-crystallin (black circles) and R54C αA-crystallin (black triangles). Black squares represent the emission spectra of bis-ANS in buffer. Grand average of hydropathicity (GRAVY) index as determined by ProtScale prediction for wild type αA-crystallin **(B)** and R54C αA-crystallin **(C)**. Red ovals depict the altered hydrophobicity in the region of the mutation (54^th^ amino acid).

### 2.3 Intracellular localization of R54C αA-crystallin

The intracellular localization of the R54C mutant and the wild type αA-crystallin was investigated using confocal fluorescence microscopy. As shown in **figure 5 (top panel)**, wild type αA-crystallin was mainly localized in the cytoplasm as suggested by a diffused cytoplasmic fluorescence signal. However, as compared to wild type αA-crystallin, the R54C mutant showed a speckled appearance in the nucleus along with a diffused staining in the cytoplasm (**figure 5, lower panel)**.

**Figure 5:**
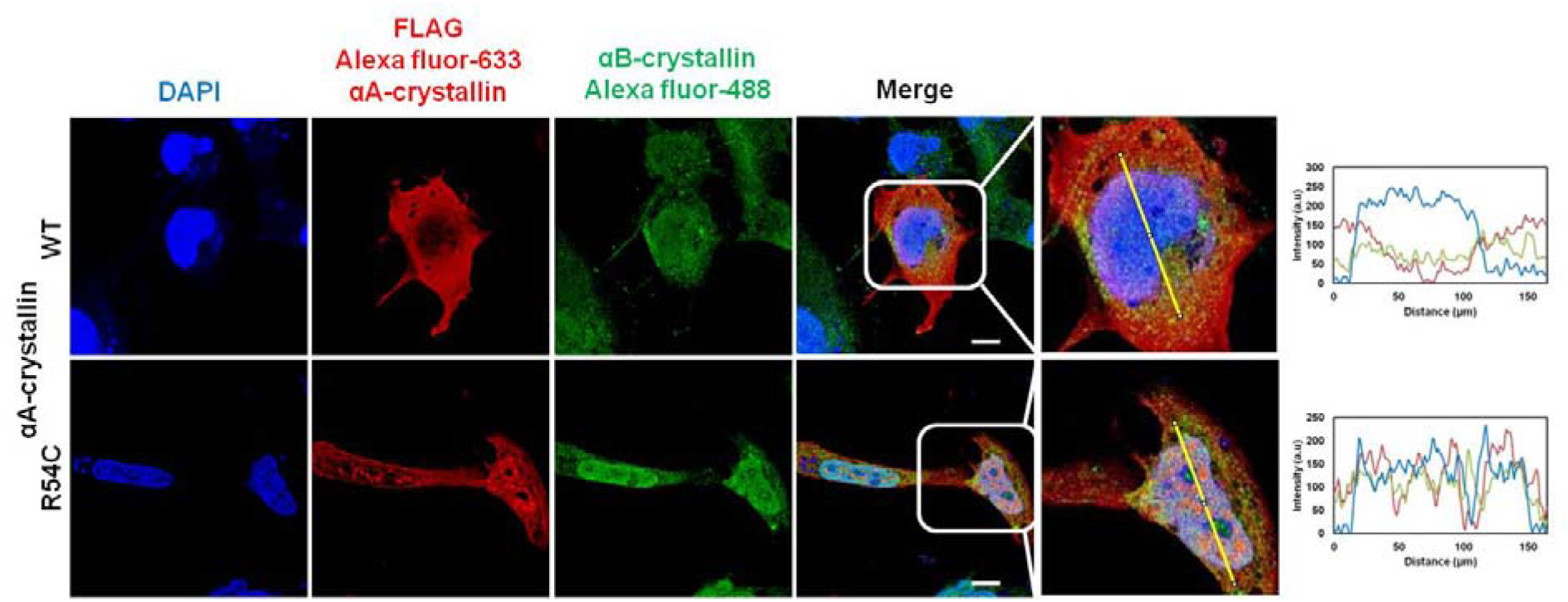
Cellular localization of wild type and R54C αA-crystallin. Confocal fluorescence microscopy of SRA 01/04 cells transfected with pCDNA3.1 vector expressing either wild type αA-crystallin or R54C αA-crystallin (red fluorescence) probed with an anti-FLAG primary antibody followed by a Alexa fluor 633 conjugated secondary antibody. The expression and localization of endogenous αB-crystallin in the transfected cells was probed using immunofluorescence (green fluorescence). (Scale bar = 10 μm). Enlarged image (right panel), of FLAG Alexa fluor-633 and αB-crystallin Alexa fluor 488 merge, shows nuclear aggregates of R54C alphaA-crystallin and enhanced αB-crystallin expression in the nucleus. Profile plots for the enlarged merge images were created using ImageJ software, depicting enhanced localization of the mutant R54C αA-crystallin in the nucleus.

We also probed the expression and localization of endogenous αB-crystallin in cells over-expressing the mutant and wild type αA-crystallin. The expression of αB-crystallin in the wild type αA-crystallin-transfected cells was found to be evenly distributed in the cytoplasm and the nucleus. However, in the cells expressing R54C αA-crystallin, the expression and localization of αB-crystallin was found to be increased in the nucleus **(figure 5, lower panel)**. Profile plots, as generated by imageJ, clearly indicate the enhanced localization of αB-crystallin in the nucleus of the R54C αA-crystallin expressing cells.

### 2.4 Comparison of the interactome of R54C αA-crystallin and wild type αA-crystallin

Although earlier reports have suggested the presence of wild type αA-crystallin in the nucleus [31], the increased nuclear localization of the R54C αA-crystallin mutant led us to investigate the interacting partners of the mutant protein. Point mutations have been shown to alter the interaction between small heat shock proteins and it has been suggested that such alteration could not only lead to absence of the original interaction, but also acquisition of deleterious interactions with new partner(s) [32, 33]. Since the mutant R54C αA-crystallin was found to localize in the nucleus of the SRA 01/04 cells, we probed the intracellular interaction between these proteins by immunoprecipitation using anti-FLAG agarose beads and performed mass spectrometry-based proteomic analysis. A relative intensity-based absolute quantification (iBAQ) of the proteomics results, shown in **table 2**, revealed major differences in the interacting partners of the wild type and mutant αA-crystallin.

**Table 2:** Interacting partners of wild type and R54C αA-crystallin:

|  | INTERACTING PARTNERS | ALPHA A<br>WT (IBAQ) | STANDARD<br>DEVIATION | ALPHA A<br>R54C (IBAQ) | STANDARD<br>DEVIATION |
| --- | --- | --- | --- | --- | --- |
| 1 | Alpha-crystallin A | 71.82 | 4.40 | 35.49 | 6.40 |
| 2 | HSP27 | 4.14 | 0.95 | 3.60 | 1.26 |
|  | **HISTONES** |  |  |  |  |
| 3 | Histone H2A | 1.11 | 0.35 | 6.18 | 2.13 |
| 4 | Histone H2B | 0.62 | 0.29 | 4.76 | 3.41 |
| 5 | Histone H3 | 0.09 | 0.16 | 0.96 | 1.66 |
| 6 | Histone H4 | 1.86 | 0.08 | 11.50 | 8.05 |
|  | **UBIQUITINYLATION** |  |  |  |  |
| 7 | Ubiquitin | 0.09 | 0.05 | 0.38 | 0.06 |
| 8 | E3 ubiquitin-protein ligase AMFR | 2.96 | 1.10 | 9.26 | 6.86 |
|  | **CYTOSKELETAL ELEMENTS** |  |  |  |  |
| 9 | Alpha Tubulin | 0.17 | 0.07 | 0.36 | 0.09 |
| 10 | Vimentin | 2.12 | 0.26 | 2.59 | 0.93 |
| 11 | Collagen alpha-1(I) chain | 0.00 | 0.00 | 0.01 | 0.01 |
|  | **TRANSCRIPTION FACTORS** |  |  |  |  |
| 12 | rRNA 2-O-methyltransferase<br>fibrillarin | 0.03 | 0.01 | 0.08 | 0.08 |
| 13 | Homeobox protein SIX3 | 2.58 | 4.46 | 0.00 | 0.00 |
| 14 | TATA-binding protein-associated<br>factor 2N | 0.00 | 0.00 | 0.04 | 0.07 |
| 15 | Nucleoside diphosphate kinase | 0.14 | 0.10 | 0.45 | 0.31 |
| 16 | 60S ribosomal protein L8 | 0.00 | 0.00 | 0.18 | 0.32 |
| 17 | Elongation factor 1-alpha 1 | 0.40 | 0.31 | 0.23 | 0.04 |
| 18 | Chromodomain-helicase-DNA- binding protein 3 | 0.00 | 0.00 | 0.04 | 0.06 |
|  | **MISCELLANEOUS** |  |  |  |  |
| 19 | Alpha-synuclein | 3.18 | 4.11 | 0.30 | 0.27 |
| 20 | Glyceraldehyde-3-phosphate dehydrogenase | 0.11 | 0.02 | 0.37 | 0.20 |
| 21 | CAS1 domain-containing protein 1 | 0.04 | 0.07 | 0.00 | 0.00 |
| 22 | Enhancer of rudimentary homolog | 0.77 | 0.56 | 2.23 | 1.47 |
| 23 | Protein LSM14 homolog A | 0.02 | 0.00 | 0.09 | 0.01 |
| 24 | Protein S100-A9 | 0.60 | 0.11 | 1.68 | 0.93 |
| 25 | Junction plakoglobin | 0.06 | 0.02 | 0.21 | 0.13 |
| 26 | Desmoplakin | 0.05 | 0.02 | 0.17 | 0.10 |
| 27 | Dermcidin | 6.63 | 1.76 | 15.66 | 8.66 |
| 28 | Heterogeneous nuclear ribonucleoprotein U | 0.13 | 0.03 | 1.07 | 0.83 |
| 29 | Desmoglein-1 | 0.15 | 0.03 | 0.47 | 0.26 |
| 30 | Uncharacterized protein C15orf52 | 0.00 | 0.00 | 0.06 | 0.10 |
| 31 | Plasminogen activator inhibitor 1 RNA-binding protein | 0.02 | 0.02 | 0.14 | 0.02 |
| 32 | Progesterone receptor | 0.00 | 0.00 | 0.79 | 1.36 |
| 33 | Lysozyme C | 0.08 | 0.15 | 0.68 | 0.62 |

Major differences were found to be in relative concentrations of histones, ubiquitinylation-related proteins and intercellular interaction proteins. The histones identified with an increased concentration upon immunoprecipitation with the mutant protein were H2A, H2B, H3 and H4, which interestingly constitute the nucleosome **(figure 6)**. Other major variations were found in ubiquitinylation-related proteins and desmosomal proteins involved in cell-cell adhesion.

**Figure 6:**
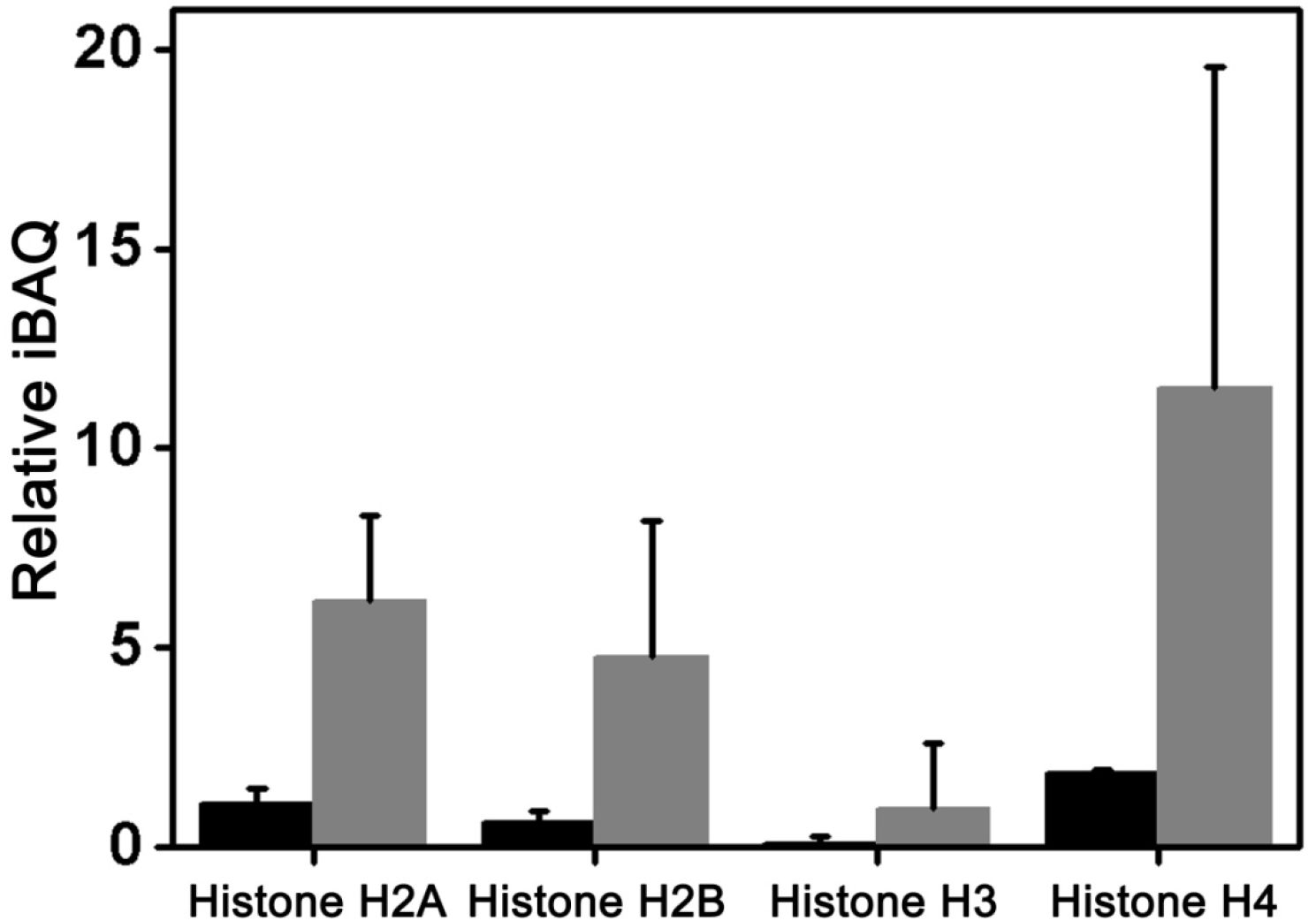
Histone binding to wild type and αA-crystallin. Relative iBAQ concentrations of different histones found to interact with wild type (black bars) and mutant R54C (grey bars) αA-crystallin as determined by immunoprecipitation followed by intensity based absolute quantification (iBAQ).

### 2.5 Intra-nuclear localization of wild type and R54C αA-crystallin

The proteomic study revealed an increased association of the mutant protein with histones. This association could be either with the nucleosome/chromatin or with the soluble pool of histones or both. We investigated whether the association of the mutant protein is with nucleosome/chromatin. We performed chromatin enrichment followed by western blotting to elucidate the binding of the mutant with DNA chromatin-associated histones. As shown in **figure 7**, the association of the mutant αA-crystallin was found to be increased with the chromatin fraction as compared to the wild type protein. The interaction of the mutant with the histones was further investigated by an immunofluorescence experiment. When probed using anti-histone H3 antibody, the mutant crystallin was found to co-localize with histone H3 **(figure 8)**. Our immunofluorescence and western blotting of the chromatin-enriched fractions suggest that the mutant protein is a part of the nucleosome complex and interacts with all four histone proteins of the nucleosome complex.

**Figure 7:**
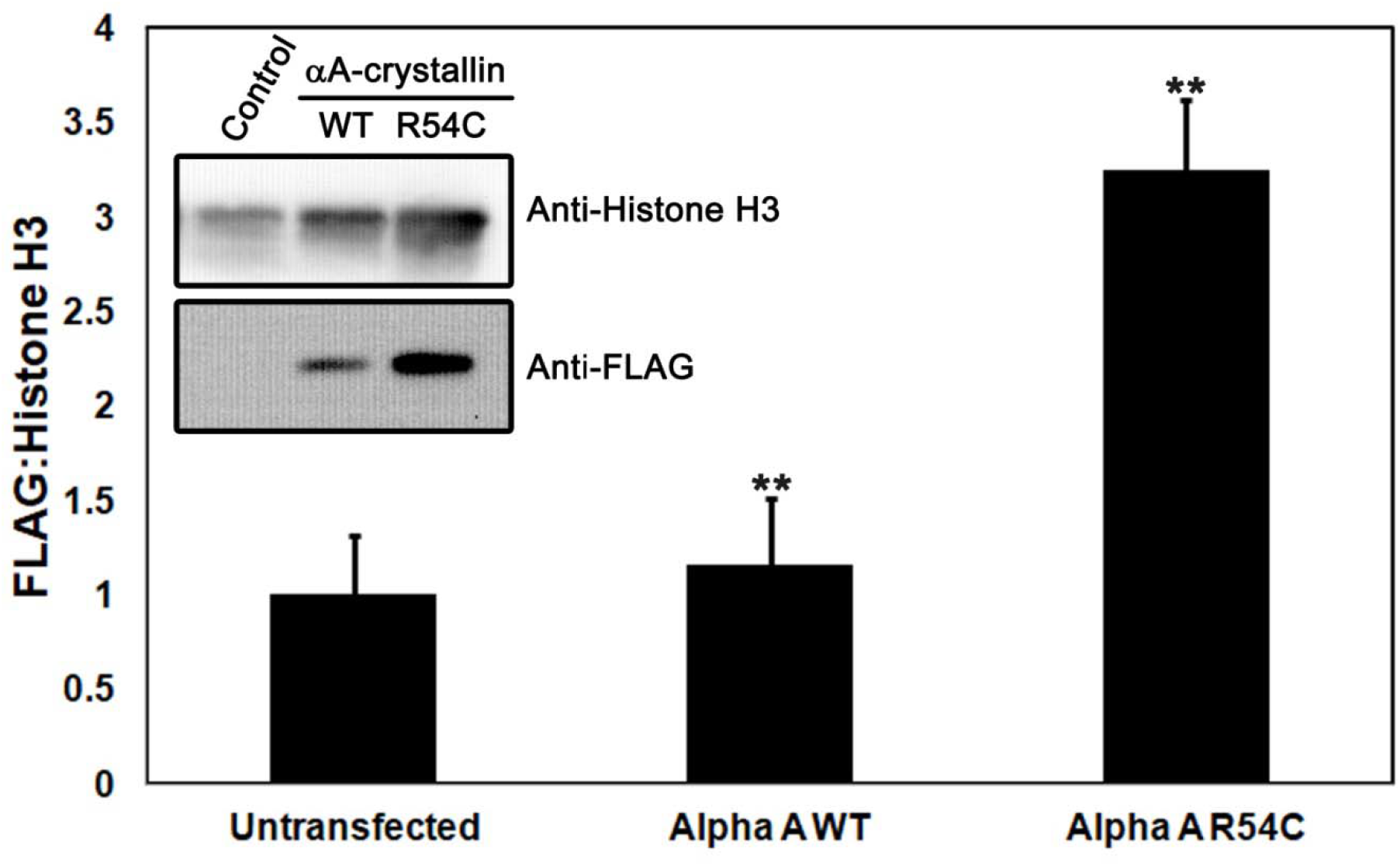
Interaction of wild type and R54C αA-crystallin with chromatin. Chromatin fractions from control cells and cells expressing wild type αA-crystallin and R54C αA-crystallin were isolated. The samples were processed and subjected to polyacrylamide gel electrophoresis as described in the materials and methods. Western blot image in the inset shows an increased level of R54C αA-crystallin in the chromatin enriched fraction as compared to that of wild type αA-crystallin. Image quantification was performed using imageJ. Bar graph represents the ratio of FLAG to Histone H3 for each group. Band intensities for FLAG:Histone H3 in the untransfected samples were normalized to 1. (n = 3; ** p < 0.01; two-tailed Welch’s t-test).

**Figure 8:**
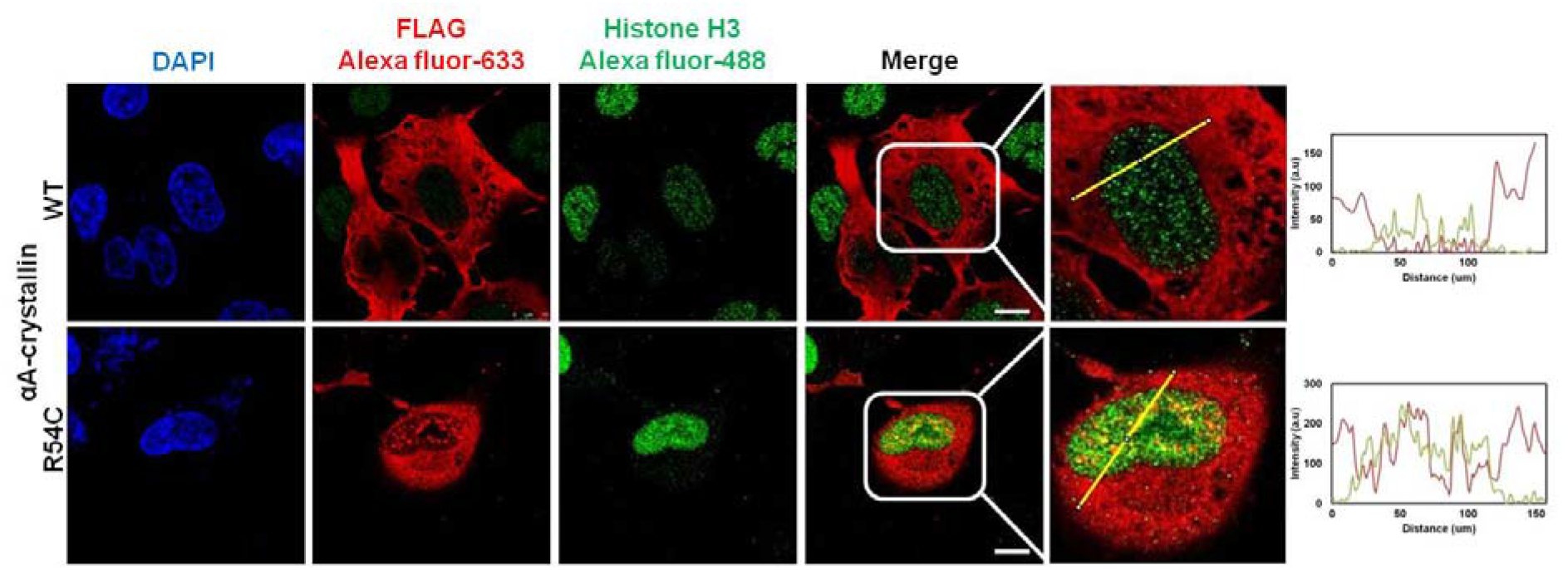
Interaction of wild type and R54C αA-crystallin with chromatin. Confocal fluorescence microscopy of SRA 01/04 cells transfected with pCDNA3.1 vector expressing either WT αA-crystallin or R54C αA-crystallin (red fluorescence) probed with an anti-FLAG primary antibody followed by Alexa fluor 633 conjugated secondary antibody. The expression and localization of histone H3 was probed using anti-histone H3 antibody followed by staining with Alexa fluor 488 (green fluorescence). (Scale bar = 10 μm). Enlarged image (right panel) of the merge of FLAG Alexa fluor-633 and Histone H3 Alexa fluor 488 shows yellow fluorescence in the case of R54C αA-crystallin but not in wildtype αA-crystallin, indicating colocalization. Profile plots for the enlarged merge images were created using ImageJ software, depicting co-localization of the mutant R54C αA-crystallin with Histone H3.

### 2.6 Effect of R54C αA-crystallin expression on cytoskeletal elements

The effect of expression of the R54C mutant and wild type αA-crystallin in SRA 01/04 cells on the organisation of cytoskeletal elements was studied using confocal fluorescence microscopy. As shown in **figure 9A and B (top panel)**, expression of wild type αA-crystallin had no particular effect on the organisation of actin and tubulin structure. However, as compared to the wild type αA-crystallin, the expression of the mutant R54C led to a depolymerisation of the actin and tubulin filaments **figure 9A and B (lower panel)**.

**Figure 9:**
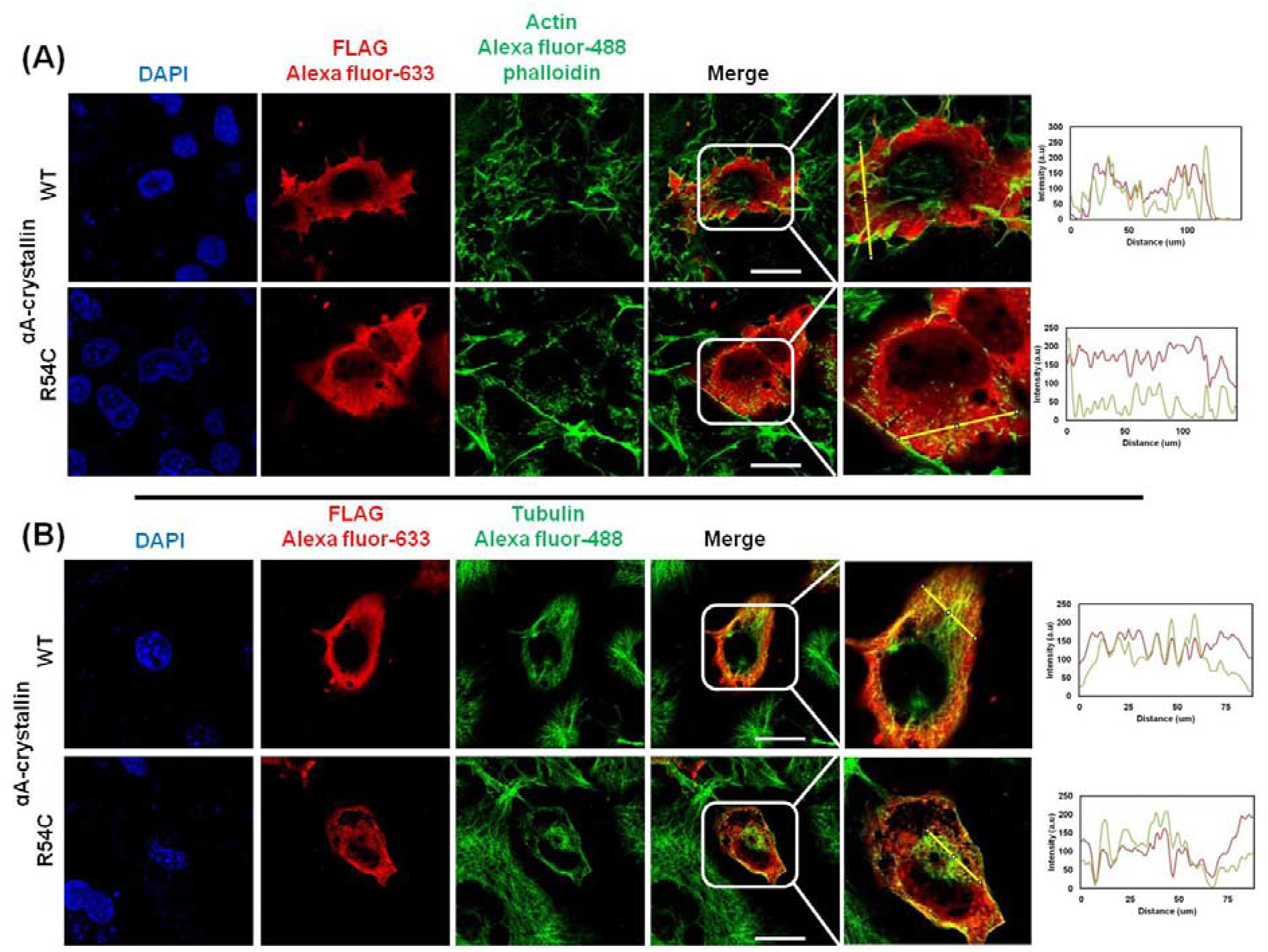
Interaction of wild type and R54C αA-crystallin with cytoskeletal elements. Confocal fluorescence microscopy of SRA 01/04 cells transfected with pCDNA3.1 vector expressing either wild type αA-crystallin or R54C αA-crystallin (red fluorescence) probed with an anti-FLAG primary antibody followed by Alexa fluor 633 conjugated secondary antibody. **(A)** The expression and localization of endogenous actin in the transfected cells was probed using Alexa fluor 488 phalloidin (green fluorescence). Profile plots for the enlarged merge images were created using ImageJ software, depicting co-localization of the wild type αA-crystallin with actin. There was minimal co-localization of the R54C αA-crystallin with actin and the actin intensity was in general low, suggesting depolymerisation of actin filaments. **(B)** The expression and localization of endogenous tubulin in the transfected cells was probed with an antitubulin primary antibody followed by Alexa fluor 488 conjugated secondary antibody (green fluorescence). Profile plots for the enlarged merge images were created using ImageJ software, depicting co-localization of the wild type and mutant R54C αA-crystallin with tubulin (Scale bar = 25 μm). Enlarged image (right panel) of the merge shows the filamentous actin Alexa fluor-488 phalloidin (9A), and tubulin Alexa fluor-488 staining (9B) for wild type alpha A-crystallin transfected cells, which is not seen in the mutant R54C alphaA-crystallin transfected cells, suggesting a stress-like response.

### 2.7 Cell death associated with R54C αA-crystallin

The effect of the expression of the mutant R54C αA-crystallin on SRA 01/04 cell survival was studied by MTT assay. The transfection efficiency of the cells was low (30%) and the selection of these cells using geneticin to create a stable transfectant pool was not possible due to cell death. MTT assay results, as shown in **figure 10A**, revealed cell death in mock-transfected cells (~34.5%) as well as in wild type αA-crystallin-transfected cells (32.85%). This could be due to the effect of the transfection reagent, lipofectamine, on the cells. Interestingly, in cells transfected with R54C αA-crystallin, cell death was found to be ~57.21%. Considering the fact that the transfection efficiency was as low as 30%, the cell death in cells transfected with R54C αA-crystallin is significantly higher than that observed in mock-transfected cells and wild type αA-crystallin-transfected cells. Further, in order to probe the mechanism of cell death, we have studied Caspase 3 activation in cells transfected with the mutant and wild type αA-crystallin. As shown in **figure 10B**, Caspase 3 was found to be activated/fragmented in cells transfected with R54C αA-crystallin. On the other hand, activation of caspase 3 was not observed in mock-transfected cells and cells transfected with wild type αA-crystallin.

**Figure 10:**
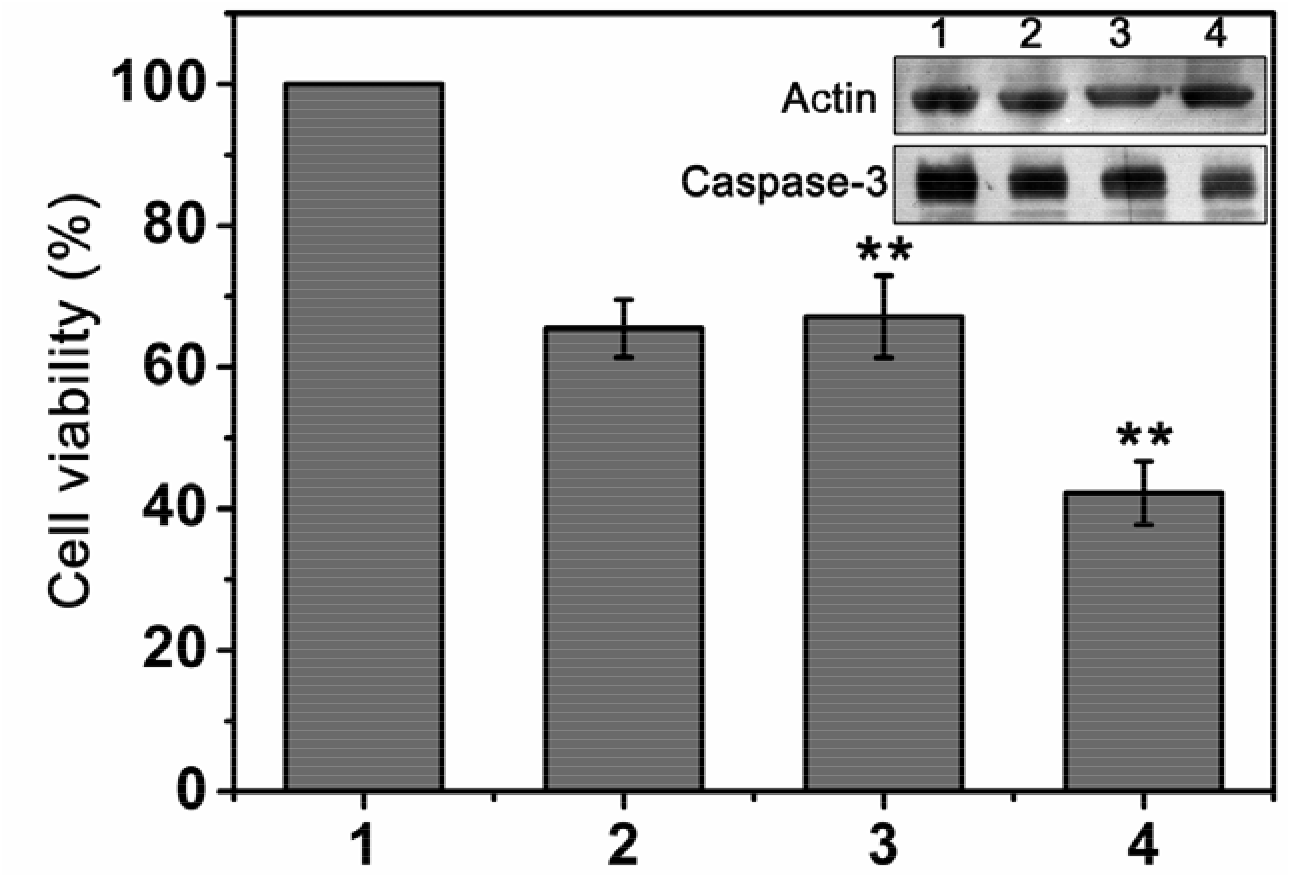
Cytotoxicity of wild type and R54C αA-crystallin. (A) MTT assay for the quantification of cell death associated with the expression of R54C αA-crystallin (n = 3; ** p < 0.01; two-tailed Welch’s t-test). Inset shows decrease in the expression of pr-caspase-3 in cells transfected with R54C αA-crystallin suggesting apoptosis (1: Untransfected cells, 2: Cells transfected with empty vector; 3; Cells transfected with wild type αA-crystallin and 4: Cells transfected with R54C αA-crystallin). (Arrow indicates fragmented caspase 3 band prominent in the mutant R54C αA-crystallin transfected cells).

## 3 Discussion

α-Crystallin is known to be involved in various cellular functions [33, 34]. The primary structure of the protein comprises of three domains *viz*. the N-terminal domain, α-crystallin domain and the C-terminal extension. Various studies have suggested the role of the N-terminal in sub-unit exchange, oligomerization and chaperone-like activity [16, 18]. Many mutations in αA-crystallin have been reported to cause cataract. Interestingly, mutations in all the conserved arginine residues (R12, R21, R49 and R54) in the N-terminal domain of the protein have been reported to cause cataract. Understanding the mechanism of cataract caused by these mutations thus becomes important.

In an attempt to gain an insight into the structural variations caused by R54C mutation in αA-crystallin, we cloned, expressed and purified the mutant R54C αA-crystallin protein and compared its structure and chaperone-like activity with wild type αA-crystallin. Our results show slight secondary and tertiary structural differences of the mutant as compared to wild type αA-crystallin. The mutant also showed an increase in the oligomer size, hydrodynamic radius and an increase in the surface hydrophobicity.

Several point mutations in αA- and αB-crystallin are known to result in the loss of chaperone-like activity, thereby causing cataract. Mutation of F71 and R116 in αA-crystallin and R120 in αB-crystallin has been shown to severely compromise the chaperone-like activity of these proteins [25, 35–37]. The G98R mutation in αA-crystallin showed a substrate-specific chaperone-like activity. While the chaperone-like activity was severely compromised for DTT-induced insulin aggregation [36], the activity was found to be slightly compromised for thermal aggregation of alcohol dehydrogenase, ovotransferrin and βB2-crystallin [38]. Interestingly, an intact or enhanced chaperone-like activity was observed against the thermal aggregation of citrate synthase [38, 39]. On the other hand, no major changes have been reported in the chaperone-like activity of the R49 and R54 mutants of αA-crystallin [25, 26]. Still other mutations *viz*. Y118 in αA-crystallin [27] and R11 [40], Q151, 450DelA and 464DelCT [41] in αB-crystallin have been reported to have an improved chaperone-like activity compared to the wild type protein. Despite the minor structural variations observed for the mutant R54C αA-crystallin, the chaperone-like activity of the mutant and wild type αA-crystallin was found to be comparable when studied against DTT-induced insulin aggregation and temperature-induced ADH aggregation (data not shown). These results are in conformity with those shown earlier for the R54C mutant of αA-crystallin [25]. Thus, factors in addition to mutational effects on the general chaperone activity of α-crystallin also seem to play a critical role in the pathology. However, these factors have not been adequately studied so far.

To gain further insights into the mechanistic details of the cataract caused by the R54C mutation, we expressed the mutant and wild type proteins in human lens epithelial cells (SRA 01/04). Immuno-fluorescence studies show that the expression of the wild type αA-crystallin is diffused across the cytoplasm. Whereas, the R54C mutant protein is seen as speckles, both in the cytoplasm and in the nucleus suggesting protein translocation and aggregation. A similar observation has been reported by Mackay et al. in the case of R49C αA-crystallin transfected cells [24]. The expression of mutant αA-crystallin affects the localization of αB-crystallin as well. In the cells expressing wild type αA-crystallin, the expression of αB-crystallin is seen to be evenly distributed both in the cytoplasm and in the nucleus. Whereas, in the cells expressing the R54C αA-crystallin, the localization is altered; there is considerably more αB-crystallin in the nucleus of these cells. Mackay et al. [24] have shown that R49CαA-crystallin aggregates and translocates to the nucleus. They suggested that since αB-crystallin forms a heteromeric complex with R49C αA-crystallin, αB-crystallin is also found to be translocated to the nucleus. It is possible that the translocation of αB-crystallin to the nucleus observed by us in the present study is also due to its hetero-oligomer formation with the R54C mutant, which aggregates and translocates to the nucleus in a manner similar to that observed in the case of the R49C mutant. However, αB-crystallin is also known to translocate into the nuclei of the cells under stress [42]. The expression of mutant proteins, its aggregation and translocation into the nucleus also cause stress to the cell, leading to stress-induced translocation of αB-crystallin into the nucleus, independent of hetero-oligomer formation with αA-crystallin. The latter explanation seems to be more reasonable as we could not identify αB-crystallin in our immunoprecipitation results (**table 3**). Raju et al. have reported a speckled appearance of R54C αA-crystallin in the cytoplasm of HeLa cells [43]. Cytoplasmic aggregation of mutant α-crystallin proteins has also been previously reported. However, such aggregations in all cases except the R49C mutant of αA-crystallin have been shown to be cytoplasmic. The mutant reported in the present study *viz*. R54C αA-crystallin, is only the second such report of a mutant aggregating and localizing in the nucleus.

To gain insights into the difference in functioning of the wild type and mutant proteins we studied the “interactome” of the two proteins. The results of the proteomic study reveal a striking increase (~10 fold) in the concentration of histones interacting with the mutant as compared to the wild type. The major histones identified in the immunoprecipitation experiments were H2A, H2B, H3 and H4; histones constituting the nucleosome [44]. Isolation and probing chromatin enriched fractions for the wild type and mutant proteins demonstrated an increased association of the R54C αA-crystallin with chromatin-enriched fractions, further confirming the association with the nucleosome. Histones are important proteins with a long half-life. Their association with wild type αA-crystallin has also been observed previously [32, 45], suggesting a possibility of protection of histones by crystallins in maintaining their cellular functions. More recently a direct interaction of αA-crystallins with histones was reported by Hamilton and Andley [46]. It was suggested that histones interact with crystallins and leads to alterations in the oligomeric status of the crystallin protein. The oligomeric changes, however, depend on the histone to crystallin ratio with a low histone to crystallin ratio resulting in water-soluble complexes while a high histone to crystallin leads to water-insoluble complexes. A further increase in histone to crystallin ratios results in the formation of both water soluble and insoluble complexes. An increase in the abundance of histones has also been reported in αA/αB double knock-out mice lenses and in lens cells expressing αA-crystallin mutants [32], probably due to an effort on the part of cells to maintain a functional pool of histones for normal cellular activities. The accumulation of histone transcripts has also been shown in lens epithelial cells upon stress *viz*. irradiation with ultraviolet C radiation waves [47]. In the study by Hamilton and Andley [46], it was speculated that the nuclear aggregation of the R49C αA-crystallin protein is a direct consequence of the increased histone to αA-crystallin ratio, resulting in water-insoluble complexes. This aggregation may sequester the histones, preventing their binding to the chromatin. In our study we show that although the mutant R54C αA-crystallin interacts with histones, the histones + R54C αA-crystallin complex still bound to chromatin. The exact consequence of this interaction still needs to be elucidated, but this may lead to an alteration in the trasncriptomic profiles, as observed in mouse lenses expressing R49CαA-crystallin [48].

The R54C mutation in αA-crystallin leads to recessive cataracts in humans. Interestingly, histones have been shown to play an important role in the regulation of Wnt signalling pathway, which is known to be important in lens development and morphogenesis and has been linked with cataract formation in different model systems [49, 50]. It can thus be speculated that the increased association of the mutant αA-crystallin with nucleosomal histones, may modulate cellular transcriptional activity including that mediated by β-catenin thereby affecting the Wnt signalling pathway. This may be a possible reason for improper lens development and cataract formation in the mutant lenses [51]. Our findings show increased interaction of various junction proteins *viz*. plakoglobin, desmoplakin and desmoglein in the lens epithelial cells. Cadherin-based junctions are important features of undifferentiated lens epithelial cells and aid in the organization of cytoskeletal elements during lens development [52]. A sequestration of these proteins by the mutant protein may thus lead to impairment in lens fibre formation and development.

α-Crystallin is known to interact with a variety of client proteins and stabilize them [33]. The role of chaperoning and stabilizing actin and tubulin by αB-crystallin has been well established [53, 54]. Also, the expression of αB-crystallin is known to increase in the presence of microtubule-destabilizing drugs, confirming its role in microtubule assembly [55]. Both αA-crystallin and actin have been shown to be important for the proper development of the lens. Further, mutant R116C αA-crystallin has been shown to exhibit decreased interaction with actin compared to wild type αA-crystallin. It has therefore been suggested that decreased interaction of mutant αA-crystallin may disturb the normal differentiation process, causing lens opacity [56]. Our work for the first time reports the destabilising nature of a mutation in αA-crystallin on the cytoskeletal elements. However, the destabilization of microtubule assembly observed with the over-expression of the R54C αA-crystallin could also be due to lack of sufficient αB-crystallin in the cytosolic compartment of the cell, due to its translocation in the nucleus.

α-Crystallins along with other sHSPs have been reported to prevent apoptosis induced by a variety of factors *viz*. oxidative stress, staurosporine, TNF-α, etc [57–59] and may exhibit this anti-apoptotic effect through various ways. For instance, αB-crystallin translocates to the mitochondria during oxidative stress and protects mitochondrial membrane potential [60]. Both αA- and αB-crystallin bind to Bax and Bcl-X(S) (pro-apoptotic molecules) and prevent their translocation to the mitochondria [61]. They are also known to suppress the activation of caspase 6 and caspase 3 and prevent apoptosis [62]. Mutations in α-crystallins are known to reduce its anti-apoptotic activity, and may even lead to induction of apoptosis [24, 63]. The activation of caspase 3 and induction of apoptosis observed in the present study in R54C αA-crystallin-transfected cells could be due to induction of apoptosis by the mutant protein and also its inability to bind and inhibit activation of caspase 3.

To summarize, the R54C mutation in αA-crystallin does not result in a significant loss of structure or chaperone activity of the αA-crystallin protein. It, however, leads to aggregation and nuclear translocation of the protein when expressed in cell culture. An altered interactome of the mutant and its association with nucleosome triggers a stresslike response, which results in a cascade of events including translocation of αB-crystallin to the nucleus, depolymerisation of the cytoskeletal elements and eventual cell death. This seems to be the plausible molecular mechanism underlying the mutation-induced cataract formation. As various neurodegenerative diseases (such as aggregate formation by polyglutamine (polyQ)) [64] and cataract [65] are characterized by the formation of nuclear inclusions of abnormal proteins, our study may also shed light on the possible mechanism of stress and cell death caused by such nuclear inclusions.

## 4 Materials and Methods

### 4.1 Materials

pET-21a (+) vector, T7 promoter and T7 terminator primers were obtained from Novagen (Madison, WI). pcDNA3.1 was obtained from Addgene (Cambridge, MA). Mutagenic primers and αA-crystallin-specific primers with restriction sites were obtained from Bioserve. Gel filtration chromatographic medium Bio-Gel A-1.5m was purchased from Bio-Rad Laboratories (Hercules, CA). Q-Sepharose matrix, Superose-6 HR 10/30 column and high molecular weight protein calibration kit comprising of blue dextran (2000 kDa), thyroglobulin (669 kDa), ferritin (440 kDa) and catalase (232 kDa) were purchased from Amersham Biosciences (Uppsala, Sweden). Insulin, dithiothreitol (DTT), Alcohol dehydrogenase (ADH), Anti-FLAG antibody and Anti-FLAG antibody-conjugated agarose beads were obtained from Sigma (St. Louis, MO). 1,1’-bi(4-anilino) naphthalenesulfonic acid (Bis-ANS) and Alexa Fluor^®^ 488 phalloidin were purchased from Thermo Fisher Scientific (Waltham, MA, USA). Antibodies against αB-crystallin, caspase 3 and actin were obtained from Abcam (Cambridge, UK). Alexa Fluor 488 and Alexa Fluor 633 conjugated secondary antibodies were obtained from Thermo Fisher Scientific (Waltham, MA, USA). HRP-conjugated secondary anti-rabbit and anti-mouse IgG antibodies were obtained from PerkinElmer (Waltham, MA, USA). Vectashield antifade mounting medium with 4’,6-diamidino-2-phenylindole (DAPI) was obtained from Vector Laboratories (Burlingame, CA, USA). 3-(4,5-dimethylthiazol-2-yl)-2,5-diphenyltetrazolium bromide (MTT) reagent was obtained from Calbiochem (San Diego, CA, USA).

### 4.2 Creating R54C mutant of αA-crystallin

pET-21a(+) vector with recombinant human αA-crystallin gene cloned between NdeI and HindIII restriction sites was used as a template to generate the R54C mutant. Briefly, the mutant was generated by mutating the 54^th^ codon, CGC to TGC, using polymerase chain reaction (PCR). Two independent polymerase chain reactions (PCRs) were performed using i) a forward T7 promoter primer and the mutagenic primer 5’-CAG CAC GGT GCT GAA GAG GGA C-3’ as the reverse primer, ii) the mutagenic primer 5’-G TCC CTC TTC TGC ACC GTG CTG-3’ as the forward primer and T7 terminator primer as the reverse primer. The resulting partially overlapping fragments were reamplified using T7 promoter and T7 terminator primers. The amplified fragment was digested and ligated in the NdeI and HindIII sites of the pET-21a(+) expression vector. The sequence of the construct was verified by T7 promoter and T7 terminator primers using 3700 ABI automated DNA sequencer.

For cloning the respective genes in mammalian expression vector, pcDNA3.1, the gene sequence of the wild type and the mutant protein were amplified using PCR with primers specific for αA-crystallin having restriction sites EcoRI and XhoI (forward primer 5’-CCG GAA TTC ATG GAC GTG ACC ATC CAG-3’ and reverse primer 5’-CCG CTC GAG TTA GGA CGA GGG AG-3’). The amplified fragment was digested and ligated in the EcoRI and XhoI sites of the pcDNA3.1 mammalian expression vector. The sequence of this construct was verified by primers specific for αA-crystallin using ABI 3700 automated DNA sequencer.

### 4.3 Expression and purification of the recombinant wild type and R54C mutant protein

For protein expression, plasmids containing the wild type and the mutant gene were transfected in *Escherichia coli* BL21 (DE3) cells. When the optical density of the culture reached 0.4 to 0.6 OD, protein expression was induced by the addition of IPTG (1 mM). Cultures were incubated for 5 h at 37 °C with stirring at 180 rpm. Cells were subsequently harvested by centrifugation at 3000 x g. The cell pellet obtained was resuspended in 50 mM Tris-HCl buffer (pH 7.2) containing 100 mM NaCl and 1 mM EDTA (TNE buffer), sonicated and centrifuged. The wild type as well as the mutant protein partitioned in the soluble fraction and were purified as described earlier [6]. Briefly, the soluble fraction was subjected to fractionation with ammonium sulphate (30-60% saturation) followed by gel filtration (Bio-Gel A-1.5m) and anion-exchange chromatography (Q-Sepharose). The concentrations of both the wild type and the mutant protein samples were estimated using its extinction coefficient (14565 M^−1^ cm^−1^ for both proteins) at 280 nm as described by Pace et al. [66].

### 4.4 Fluorescence studies

The tryptophan fluorescence spectra of wild type and R54C αA-crystallin were monitored in TNE buffer with a protein concentration of 0.2 mg ml^−1^. Spectra were recorded using an excitation wavelength of 295 nm with excitation and emission band passes set at 2.5 nm each. Emission spectrum was recorded from 300 nm to 400 nm.

The surface hydrophobicity of the wild type and mutant proteins was probed by recording the fluorescence spectra of the hydrophobic probe, bis-ANS (10 μM), as reported previously [67], upon incubation with 0.2 mg ml^−1^ of the respective protein samples at 25 °C for 10 min. An excitation wavelength of 390 nm was used with excitation and emission band passes set at 2.5 nm each. Fluorescence spectra were recorded from 400 to 600 nm. All spectra were recorded in the corrected spectrum mode using a Hitachi F-4500 fluorescence spectrophotometer.

### 4.5 Local hydrophobicity prediction

The hydrophobicity values around the wild type and mutated amino acid were predicted using the ProtScale online software at ExPASY server (http://web.expasy.org/protscale) with the Kyte-Doolittle hydrophobicity scale.

### 4.6 Circular dichroism studies

Circular dichroism spectra of the proteins were recorded as described previously [68]. Briefly, a concentration of 1.0 mg ml^−1^ of protein was used in TNE buffer in a 1 cm path length cell for near-UV region and a concentration of 0.2 mg ml^−1^ of protein in a 0.1 cm path length cell for the far-UV region. All spectra were recorded using a JASCO J-715 spectropolarimeter at 25 °C. The spectra reported are the average of 4 accumulations. Appropriate spectra of buffer blanks run under the same conditions were subtracted from the sample spectra and the observed ellipticity values were converted to mean residue ellipticity (MRE). CD data analysis was performed by using the CAPITO (**C**D **A**nalysis and **P**lott**i**ng **To**ol) CD analysis online software, (Leibniz Institute on Aging – Fritz Lipmann Institute) [28].

### 4.7 Size exclusion chromatography

The oligomeric sizes of the wild type and mutant proteins were evaluated using a Superose-6 HR 10/30 pre-packed FPLC column (dimensions 1×30 cm). The proteins thyroglobulin (669 kDa), ferritin (440 kDa) and catalase (232 kDa) were used as high molecular mass standards.

### 4.8 Dynamic light scattering studies

For DLS studies, protein samples (2 mg ml^−1^) were centrifuged and filtered through a 0.22 μ membrane filter. Measurements were carried out using a Photocor Dynamic Light Scattering Instrument (Photocor Instruments Inc., MD) equipped with a 633 nm 25 mW laser. All measurements were performed at 90° angle with temperature set at 25 °C. Data was analysed using the Dynals (version 2.0) software provided with the instrument.

### 4.9 Intracellular localization of wild type αA-crystallin and R54C mutant

Human lens epithelial cells (SRA 01/04) were cultured in DMEM, supplemented with 10% FBS and antibiotics (5 μg ml^−1^ penicillin & 6 μg ml^−1^ streptomycin) at 37 °C under 95% humidity and 5% CO_2_. For transfection studies, SRA 01/04 cells grown on cover-slips were transfected with 1 μg of pcDNA3.1-N FLAG containing the wild type and mutant αA-crystallin using Lipofectamine 2000 reagent following the manufacturer’s protocol.

Cells were fixed 48 h post transfection with 4% formaldehyde. The fixed cells were permeabilized with 0.05% Triton X-100 for 8 min. After blocking with 2% BSA, cells were incubated for 2 h with the desired primary antibodies against FLAG tag, αB-crystallin, actin or Alexa Fluor^®^ 488 phalloidin (for staining actin filaments). Cells were then washed with PBS and further incubated with the corresponding secondary antibodies. Cells were counterstained with DAPI, and images were acquired using a 63x objective lens on a Leica confocal microscope (TCS-SP8; Leica Microsystems, Wetzlar, Germany). Images were analysed by Leica Application Suite AF software provided by the company.

### 4.10 Identification of interacting partners of wild type and mutant αA-crystallin

Cells were harvested 48 h post transfection by gentle scraping and whole cell protein was extracted by incubating cells in RIPA lysis buffer (50 mM Tris–Cl, pH-7.4, 150 mM NaCl, 1% NP-40, 0.5% sodium deoxycholate, 0.1% SDS) supplemented with 1 mM EDTA, 1 mM PMSF and 1X protease inhibitor cocktail for 30 min at 4 °C. The lysate was centrifuged at 10,000 x g for 25 min at 4 °C. The supernatant was pre-cleared by incubation with Protein G Sepharose beads for 1 h on a rotator at 4 °C. After removing the Protein G Sepharose beads by centrifugation, the protein in the supernatant was estimated by Bradford’s method according to the instructions of the manufacturer. Equal amounts of lysates for the two samples were further mixed with anti-FLAG antibody conjugated agarose beads and incubated overnight at 4 °C. Subsequently, the beads were washed thrice with Tris-buffered saline (TBS) and the proteins were eluted by adding Glycine-HCl (pH. 2.5). The eluted proteins were mixed with Laemmli buffer and loaded on a 4-12% precast Invitrogen Novex SDS gel followed by fixing and staining with Coomassie blue.

The preparation of gel slices followed by reduction, alkylation and in-gel protein digestion was done as described by Shevchenko et al.[69] The peptides were then desalted and enriched as described by Rappsilber et al.[70] Peptides eluted from desalting tips were dissolved in 2% (v/v) formic acid and sonicated for 5 min. Samples were analysed on a nanoflow LC system (Easy nLC II, Thermo Scientific) coupled to a Q-exactive mass spectrometer (Thermo Scientific). Peptides were separated on Bio Basic C18 pico-Frit nanocapillary column (75 μm × 10 cm; New objective, MA, USA) using a stepwise 60min gradient between buffer A (5% ACN containing 0.2% formic acid) and buffer B (95% ACN containing 0.2% formic acid). The HPLC flow rate was set to 400 nl per min during analysis. MS/MS analysis was performed with standard settings using cycles of 1 high resolution (70000 FWHM) MS scan (from *m/z* 400–1600) followed by 10 MS/MS scans of the 10 most intense ions with charge states of 2 or higher. Protein identification and intensity based absolute quantification (iBAQ) [71] was performed using MaxQuant (version 1.3.0.5) [72] using default settings. The human sequences of UNIPROT (version 2016-03) were used as database for protein identification. MaxQuant used a decoy version of the specified UNIPROT database to adjust the false discovery rates for proteins and peptides below 1%. Each experiment was repeated three times. Enrichment or depletion of an identified protein in at least two repeat experiments indicated alteration of abundance of the protein during the experiment.

### 4.11 Interaction of wild type and mutant αA-crystallin with chromatin

Cells were harvested 48 h post transfection as described by Kustatscher et al. [73], Briefly, cells were washed with PBS once 48 h post-transfection followed by treatment with pre-warmed (37 °C) formaldehyde (1%) for 10 min at 37 °C. Glycine (0.25 M) was used to quench the reaction followed by cell collection in PBS. The cells were lysed in lysis buffer (25 mM Tris (pH 7.4), 0.1% Triton X-100 and 85 mM KCl supplemented with 1X protease inhibitor cocktail). The nuclei were separated by centrifugation at 2300 x g for 5 min. The supernatant at this step was used to quantify the nuclei. The nuclei were then resuspended in lysis buffer and equal amounts of samples (based on the protein estimates in the supernatant) were treated with RNase A followed by centrifugation at 2300 x g for 10 min. The pellet was resuspended in SDS buffer (50 mM Tris (pH 7.4), 10 mM EDTA and 4% SDS supplemented with 1X protease inhibitor cocktail) followed by the addition of urea buffer (10 mM Tris (pH 7.4), 1 mM EDTA and 8 M urea). The contents of the tubes were mixed thoroughly followed by centrifugation at 16,100 x g for 30 min. The washing step (with SDS and urea buffer) was repeated once to wash contaminants followed by washing with just SDS buffer to remove urea. The final pellet was resuspended in storage buffer (10 mM Tris (pH 7.4), 1 mM EDTA, 25 mM NaCl and 10% glycerol supplemented with 1X protease inhibitor cocktail). The samples were then sonicated for 15 min (30 sec on & off pulse) followed by centrifugation at 16,100 x g for 30 min. The supernatant was transferred to fresh tubes containing Laemmli buffer, boiled for 10 min and resolved on a 12% SDS PAGE for western blotting as described previously [74]. Briefly, the proteins, after resolving on 12% SDS PAGE were thereafter transferred to nitrocellulose membrane (Hybond C-extra) using a semi-dry transfer apparatus. The membrane was then placed in a blocking solution (5% BSA in Tris Buffered Saline (TBS)) for 2 h followed by incubation with primary antibody (diluted in 1% BSA in TBS) followed by washing (thrice) and incubation in Horseradish peroxidase (HRP) conjugated secondary antibody. The membrane was finally washed with TBS containing 0.1% Tween-20 (TBST) thrice and developed by Vilber-Lourmat Chemiluminescence Imaging System (MArne-la-Valée Cedex 3, France) using the Chemi-Capt software, after adding the HRP substrate.

For immunofluorescence assay, transfected cells were fixed 48 h post transfection with 4% formaldehyde. The fixed cells were stained with the desired primary antibodies against FLAG tag and histone H3 followed by the corresponding secondary antibodies. Cells were counterstained with DAPI, and images were acquired using a 63x objective lens on a Leica confocal microscope (TCS-SP8; Leica Microsystems, Wetzlar, Germany). Images were analyzed by Leica Application Suite AF software provided by the company.

### 4.12 Cell death induced by R54C mutant of αA-crystallin

Cells were harvested 48 h post transfection by gentle scraping and whole cell protein was extracted by incubating cells in lysis buffer (50 mM Tris Cl, pH 8.0, containing 1% Triton X-100, 1% sodium deoxycholate, 0.1% sodium dodecyl sulphate, 150 mM sodium chloride, 1X protease inhibitor, 1X phosphatase inhibitor and 1 mM sodium orthovanadate) for 30 minutes on a rotator at 4 °C. After a brief sonication, the lysate was centrifuged (12,000 x g for 20 minutes at 4 °C) to remove any aggregates. The protein in the supernatant was estimated by Bradford’s method according to the instructions of the manufacturer. 25 μg of the total cell protein was loaded on to a 12% SDS polyacrylamide electrophoresis gel. A similar protocol as mentioned in section 2.11 was used for western blotting using anti-caspase 3 antibody.

For MTT assay, cells were grown in 96 well plates and transfected accordingly. Cells were washed with PBS 48 h post transfection and incubated with MTT reagent dissolved in DMEM. The solution was removed after 4 h and the formazan crystals formed were dissolved in isopropanol. Absorbance at 570 nm was monitored using a 96 well plate reader.

## Acknowledgements

SMA acknowledges Council for Scientific and Industrial Research (CSIR) for financial support. The authors thank Ms Shivali Rawat and Dr. Swasti Raychaudhuri for help and suggestions in proteomics data analysis. The authors also thank the proteomics facility of CCMB for the analysis. CMR acknowledges funding from the Sir J.C. Bose National Fellowship of the Dept. of Science and Technology, India.

## Author contributions

**Ch. Mohan Rao, Tangirala Ramakrishna, Bakthisaran Raman, and Saad M. Ahsan:** Conceptualization, methodology and data analysis. **Saad M. Ahsan:** Experimentation and data acquisition. **Ch. Mohan Rao, Tangirala Ramakrishna, Bakthisaran Raman, and Saad M. Ahsan:** Writing, reviewing and editing. **Ch. Mohan Rao:** Supervision and funding acquisition.

